# Concurrent maintenance of visual imagery and short-term memory provides evidence for their distinct representations

**DOI:** 10.1101/2024.05.07.593049

**Authors:** Elyana Saad, Juha Silvanto

## Abstract

Recent research indicates there is overlap in the neural resources used during imagery and visual short-term memory. But do visual short-term memory and visual imagery operate on similar representations during recall? Here we investigated this question by asking participants to perform a delayed match to sample task for the contrast of visual gratings as cues. In the “Imagery” condition, participants were asked to form an accurate mental image of the visual cue, and at the end of the trial, perform a matching task on the mental image contrast. In the “Memory” condition, participants were not required to perform visual imagery but merely instructed to perform the delayed contrast-matching task. In Experiment 1, participants were told at the beginning of each block whether to engage in memory or imagery. The results showed that for the relevant contrast feature, matching judgments were more accurately in the “Memory” than in the “Imagery” condition. Thus imagery did not maintain an accurate representation of the encoded image, even when the visual features could still be maintained in visual short-term memory. In Experiment 2, participants were required to engage in memory and imagery simultaneously, and were told which to base their judgment on after each trial. The key finding was that the superior accuracy for memory over imagery remained, indicating that the contents of VSTM and imagery are based on distinct representations.

## Introduction

The relationship between mental imagery and visual short-term memory (VSTM) has been the subject of much debate. It is generally thought that imagery and VSTM make use of the same neuronal resources in the early visual areas (see e.g. Pearson et al, 2015, Pearson & Kosslyn, 2015). This view is also supported by demonstrations of imagery affecting the encoding of incoming visual input (Pearson et al, 2008), and distracting visual information interfering with consciously experienced memory content (Perky, 1918; Bona et al, 2013). However, the relationship between conscious experience of memory content (e.g. in form of imagery) and the actual short-term maintenance of information might not be as direct as traditionally thought (Soto & Silvanto, 2014). There is both behavioral and transcranial magnetic stimulation evidence to suggest that the two might be based on distinct representations (Bona et al 2013, Saad et al, 2015, Jacobs & Silvanto, 2015).

Here we examined this issue further by comparing the fidelity of information maintenance with or without visual imagery. Some findings suggest that strong visualisation of a stimulus through mental imagery enhances maintenance of visual information (e.g. Marks, 1973), and that individuals with high imagery capability perform better in some visual memory tasks relative to low imagers (Keogh & Pearson, 2011). However, disruptive effects of mental imagery on memory performance have also been reported. When participants were asked to match a perceived color to a familiar object color from long-term memory, high imagers had worse performance than low imagers (Reisberg et al., 1986). Similar effects have been observed for the recall of the visual appearance of objects and famous faces from short-term memory (Reisberg & Leak, 1987). This imagery-associated cost in memory performance has been explained in terms of selective retrieval of visual features that are a mixture of the percept and the internally represented features (Heuer et al., 1986) hence impairing objective sensitivity. The patterns of inter-individual correlations between memory performance and the observer’s ability to engage in imagery (i.e. high *vs* low imagers) suggest that visual short-term memory and visual imagery may operate on distinct representations.

The two experiments in the present study assessed the fidelity of information maintenance with or without visual imagery. The logic was that if both “Imagery” and “Memory” conditions rely on the same representation, then the performance between them should not differ. In Experiment 1, in the “Imagery” blocks, participants were asked to form an accurate mental image of the visual cue, and at the end of the trial, match the contrast of their mental image to exemplars of different test stimuli (i.e. to indicate which of the test stimuli matched the contrast of their mental image). In the “Memory” blocks, participants were simply asked at the end of the trial to match the contrast of the initial memory cue with the contrast of the target gratings. In separate blocks, a further control condition was added requiring participants to additionally image or maintain in short-term memory the orientation of the grating in addition to its contrast. This was done to ensure that the cognitive load is constant across conditions - as a mental image is likely to contain both orientation and contrast, which are the main features of the grating.

In Experiment 2, the “Imagery” and “Memory” conditions were not blocked and hence participants did not know whether they would be required to perform the matching contrast judgment based on their own visual imagery or based merely on a memory trace. Observers were cued at the end of the trial as to whether the recall of information for the matching judgment was to occur from a imagery representation or from memory.

## Materials and Methods

### Participants

30 participants *(9 female; mean age 23 years old*) with normal or corrected to normal vision participated in the three experiments. All participants were naïve to the aim of the study, provided written informed consent before the experiment, and were monetary rewarded. The study was performed in agreement with the Declaration of Helsinki and approved by the ethics committee of the Hospital District of Helsinki and Uusimaa. All participants benefited from training blocks prior to the experiment and it was ensured that they understood and were following task the instructions when the experiment commenced.

All had taken part in prior studies on visual imagery (e.g. Saad et al, 2015, Saad & Silvanto, 2013) and thus had experience in generating mental images in an experimental setting.

### Stimuli

Stimuli and task were controlled by E-prime v2.0 (Psychology Software Tools Inc., Pittsburgh, USA; http://www.pstnet.com/eprime.cfm). All stimuli were sinusoidal luminance-modulated gratings (with a diameter of 5 degrees of visual angle; generated with custom-made software in Matlab), presented foveally from a viewing distance of 57 cm on a gray background; they were adapted from our prior studies (Silvanto & Soto, 2012, Soto et al, 2011, Saad & Silvanto, 2013; Saad et al, 2015). The spatial frequency of the gratings was 1.44 cycles/degree. The visual cues had a Michelson contrast of 0.2, 0.3, 0.4 and 0.5 whereas the probe cue was 0.2 and the contrast scale exemplars were 0.1, 0.2, 0.3, 0.4, 0.5 and 0.6. The mask was a uniformly black circle with the same diameter as the gratings. The stimuli were presented on a 22-inch screen with 1280 x1024 pixel resolution.

### General procedure

Each trial began with a 1 sec foveally fixation point, followed by the visual cue (300 ms). The contrast of the visual cue was either 0.20, 0.30, 0.40, or 0.50. In the condition *not* requiring orientation maintenance, the visual cue was always vertical; in conditions requiring orientation maintenance, its orientation was either +/-20, 30, 40 or 50 degree from the vertical. To avoid any afterimage induction by this cue, a mask (a uniformly black circle, appeared after the offset of the visual cue for 100ms). In the No Imagery condition, participants were instructed to hold the visual cue in memory; in the Imagery condition, to form an accurate mental image of the visual cue. This was followed by a delay period of either 3, 5, or 7 sec. The imagery/memory contrast was then assessed by asking participants to match it to exemplars presented at the end of the trial. Specifically, participants were shown a display with 6 gratings and asked to choose the closest match to the contrast of their memory/mental image. The Michelson contrasts of these stimuli were: 0.10, 0.20, 0.30, 0.40, 0.50 and, 0.60 and the responses on the keyboard associated with these contrasts were 1, 2, 3, 4, 5, and 6, respectively. (For example, participants pressed “1” if the contrast of imagery/memory matched with the visual cue, the contrast of which was 0.1). In the blocks where cue orientation was addressed, a probe cue (a grating tilted either +/-10 degrees relative to the visual cue) was presented after the assessment of the contrast of memory/mental image. Participants were requested to judge the direction of the tilt relatively to that of the visual cue, corresponding to the memory or the imagery conditions (i.e. tilted to the left or right).

### Experimental sessions

Two experiments were carried out (N= 10). Each experiment was run as a within-groups factor.

#### In Experiment 1

“Imagery” and “Memory” conditions were run separately (See Fig. 1A). 8 blocks of 48 trials were run for each condition. Each block contained 8 trials with each of 6 cue contrasts. In “Imagery” blocks, participants received the following instructions: “Please make a mental picture of the visual cue and hold it actively in your mind’s eye, Focus on it throughout the trial until asked to report its contrast and/or orientation.” In the “Memory” blocks, participants were instructed: “Please hold the visual cue in memory until asked to report its contrast and/or orientation.” Half of the blocks for each condition also assessed memory for the grating orientation. After the training blocks, prior to the commencement of the experiment, the experimenter discussed the instructions with the participant to ensure that they were followed.

**Figure 1.**
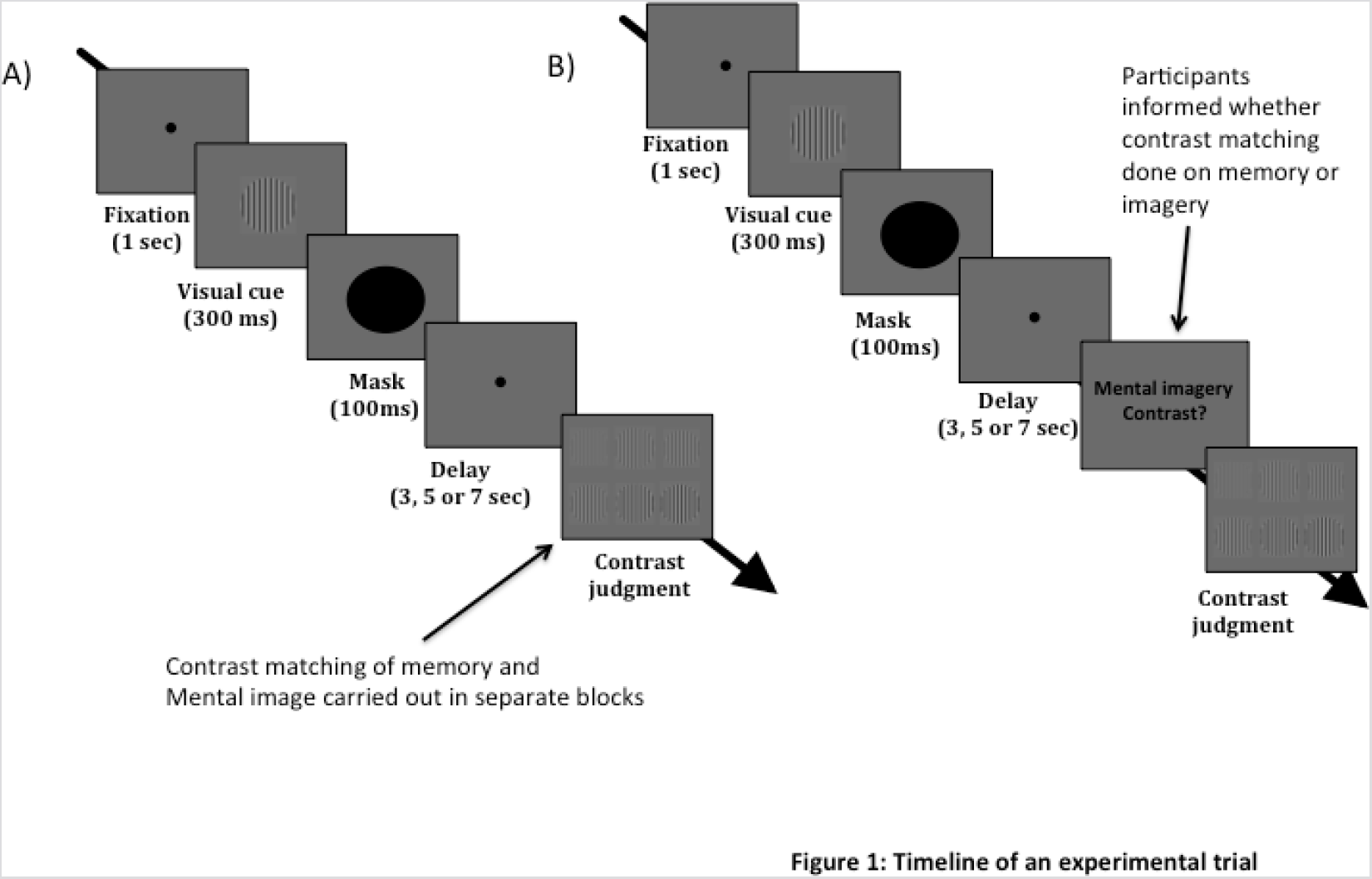
Timeline of an experimental trial. At the start of each trial, participants were presented with a visual cue grating. The task involved maintaining the contrast of the grating; in half of the blocks, orientation also needed to be maintained (when orientation was not maintained, all the gratings were vertical in orientation). Participants were instructed to carry the maintenance of the visual cue either through imagery or memory (see Methods for specific instructions). After the maintenance period, participants were asked to choose the grating that had the same contrast as their imagery/memory content. In half of the blocks, participants were additionally asked (after the contrast judgment) to perform an orientation discrimination judgment based on the memory/imagery of the visual cue; this involved indicating whether a probe cue was tilted to the left or right relative to the visual cue. In Experiment 1 figure 1A, the contrast judgment assessed selectively either mental image or memory contrast. In experiment 2 figure 1B, the assessment of the contrast judgment was intermixed within same blocks.

#### In Experiment 2

“Imagery” and “Memory” conditions were assessed on a trial-by-trial basis within the same blocks (see Fig.1 B). In other words, trials assessing Imagery contrast and Memory contrast were randomly intermixed across trials. Here participants were informed at the end of the trial whether they would need to perform the contrast matching judgment based on the imagery content or based on a mere memory trace. The instructions were as follows: “You will see a visual cue at the beginning of the trial. At the end of the trial you will be asked either to judge your memory of the visual cue or to report your imagery, based on its vividness. Thus, you need to retain a memory of the visual cue and also to make an internal mental image of it in the focus of your mind’s eye.”

8 blocks of 96 trials were run; in each block, half of the trials were for “Imagery content” and the other half assessed “Memory content”. Half of the 8 blocks also assessed memory for the grating orientation. Participants were informed in both experiments before each block whether the task involved contrast only, or both contrast and orientation to equate feature load across conditions. In order to assess contrast discrimination at baseline, (without any imagery/memory demand) a control experiment (*Experiment 3*) was run. In this experiment, the delay period between the visual cue and recall was 500ms; the timeline of the trial was otherwise identical to that in Experiment 1. Orientation discrimination was not assessed. This was run in two blocks of 64 trials. One participant was removed due to discrimination accuracy of more than two standard deviations below the group mean.

## Results

### Experiment 1

In Experiment 1, contrast of “imagery content” and “memory content” were assessed in different blocks. The accuracy of contrast matching is shown in Figure 2A. Chance level is 16.7%. Data are shown separately for the condition in which grating orientation also needed to be maintained. A repeated measure 2x2x3 ANOVA with main factors: cognitive load (contrast only; contrast and orientation); contrast matching response (imagery content; memory content), and maintenance duration (3, 5, and 7 sec) was carried out. This revealed a significant main effect of matching response (F (1,9) = 17.9 p=0.002; partial η2=0.665) but no significant effect of memory load (F(1,9)=2.41 p = 0.16; partial η2=0.211) or delay duration (F(2,18)=1.402 p = 0.27; partial η2=0.135). None of the interactions were significant. Thus the contrast matching accuracy was significantly higher in the Memory condition than in the Imagery condition. Memory accuracies for orientation information were 77% (SD: 0.09) for imagery and 78% (SD: 0.08) for VSTM, respectively.

**Figure 2.**
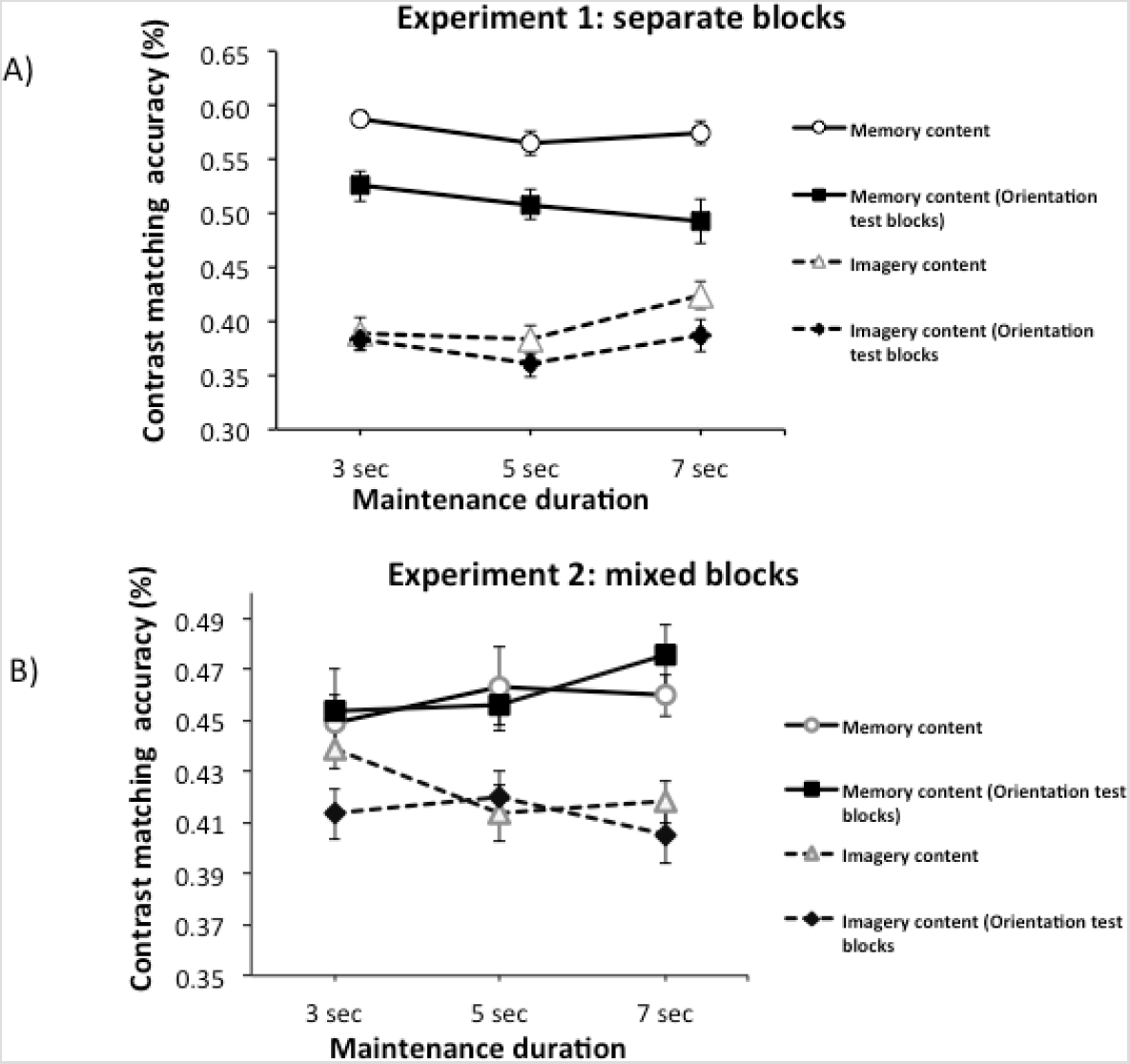
Mean accuracy contrast judgments based on imagery/memory content in the two experiments (n=10 in both). In Experiment 1 (panel A), imagery and memory content were assessed in separate blocks. Participants were either assessed for cue contrast only (“memory content” and “imagery content” conditions), or also for orientation in addition to contrast “memory content (Orientation test blocks)” and “Imagery content (Orientation test blocks)” conditions. In Experiment 2 (panel B), trials types were randomly intermixed within each block. Accuracy was higher for judgments of memory content than imagery content. Interestingly, orientation judgment accuracy did not differ between memory and imagery conditions (see Results section); thus this effect was specific to the contrast results. The Error bars indicate SDs from which between-subjects variance has been removed (Loftus & Masson, 1994).

### Experiment 2

In Experiment 2, contrast of “imagery content” and “memory content” were assessed in the same block. The accuracy of contrast matching is shown in Figure 2B. Data are shown separately for the condition in which orientation information also needed to be maintained. A repeated measure 2x2x3 ANOVA with main factors: cognitive load (contrast only; contrast and orientation); contrast matching response (imagery content; memory content), and maintenance duration (3, 5, and 7 sec) was carried out. This revealed a significant main effect of matching response (F (1,9) = 5.68 p=0.04; partial η2=0.387), but no significant effect of memory load (F(1,9)=0.019 p = 0.9; partial η2=0.002) or delay duration (F(2,18)=0.008 p = 0.9; partial η2=0.001). None of the interactions were significant.

Thus, as in Experiment 1, the contrast matching accuracy was significantly higher for the Memory condition relative to the Imagery condition. The memory accuracy for orientation information did not differ between Imagery and Memory conditions (75%; SD=0.08 for both).

Figure 3 shows the relationship between the chosen contrast at probe and the contrast of the visual cue. The dashed lines represent trials with correct responses. In both imagery and memory test conditions participants overestimated the contrast of the visual cues when these were low in contrast and underestimated the contrast of the visual cues when these were of higher contrasts. In the control experiment (Experiment 3) without any delay period, such under/overestimation did not occur. In the control experiment the accuracy of the contrast judgment was 58 % (SD: 0.1), which is similar to the level of accuracy in Experiment 1 for the memory contrast judgments. Also, the accuracy of the maintained information did not decay during the delay period.

**Figure 3.**
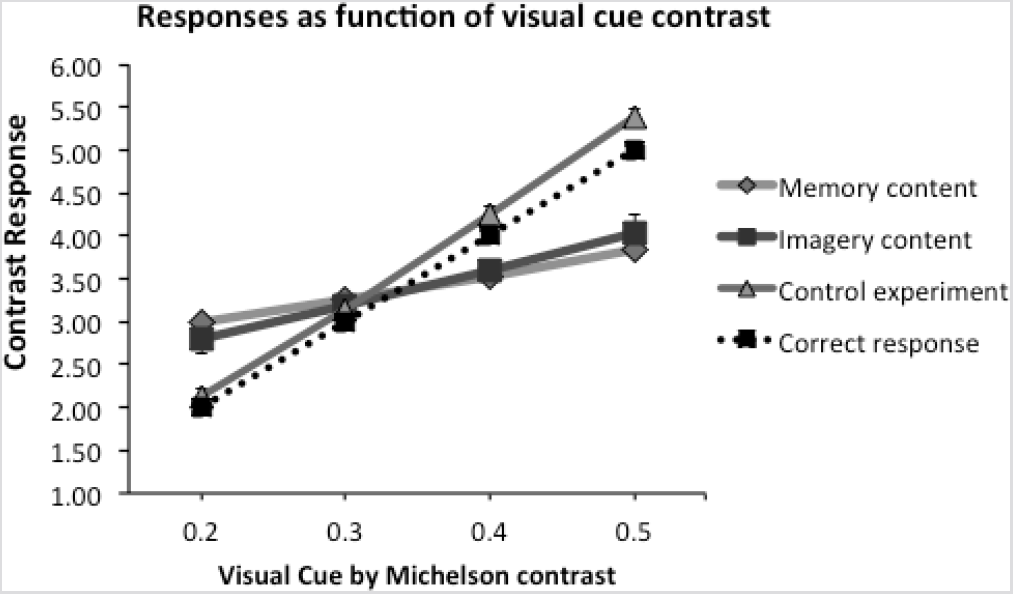
Mean contrast judgments as a function of the contrast of the visual cue. Mean contrast judgments as a function of the visual cue contrast. For imagery and memory conditions, the data across the two experiments was combined. The dashed line indicates the correct response (i.e. for cue contrast of 0.2, the correct match on the scale was 2; for cue contrast 0.3, the correct match on the scale was 3, and so on). The figure shows that, for both memory and imagery, participants overestimate the contrast of visual cue with low contrast, and underestimate them for high contrasts. In the control experiment with a short delay (500 ms) not requiring memory maintenance, such under/overestimation did not occur. The Error bars indicate SDs from which between-subjects variance has been removed (Loftus & Masson, 1994).

## Discussion

Our results show that contrast judgments were matched with a higher fidelity to the contrast of the visual cue when the participants were instructed to base their judgments on a memory representation of the visual cue relative to a context in which participants used visual imagery contents to perform the matching task. Specifically, Experiment 1 showed that when participants were asked to form a mental image of the visual cue and use the mental image contrast for the matching test, the accuracy of the contrast judgment was significantly lower compared to conditions with no imagery requirement and where participants were asked to report the contrast of the memory content. In Experiment 2, trials assessing “Imagery contrast” and “Memory contrast” were intermixed within blocks and whether participants had to use a memory of the visual cue or their visual imagery was only indicated at the end of the trial. Thus participants were involved in both mental imagery and memory maintenance simultaneously (which was not the case in Experiment 1). Critically, the pattern of results held across Experiment 1 and 2 regardless of whether participants knew in advance if they would need to use their imagery or not, hence ruling out that the reported effects could be due to differences in the way that the visual cues were encoded. These effects were not modulated by the duration of the maintenance period suggesting that the origin of the effects is not perceptual in nature but makes use of representations that are sustained over the delay period (Magnussen et al., 2003). Our results also show that both mental imagery and memory were associated with an overestimation of the contrast of visual cue with low contrast cues, and an underestimation for high contrasts (see Figure 3). In the control experiment without delay period (where maintenance was thus not required), such under/overestimation did not occur.

It is important to note that these results were obtained with a method for assessing memory/imagery contrast which was likely to rely strongly on the subjective experience of that content. Specifically, the task required matching an internal image to an external stimulus; this is likely to require a level of introspection not needed to perform a simple forced-choice discrimination task. In this study, orientation maintenance was assessed using a forced-choice task; this may explain why performance did not differ between imagery and memory. Furthermore, our prior studies using forced-choice tasks have found similar levels of performance for imagery and memory (e.g. Saad & Silvanto, 2013; Saad et al, 2015). Thus it appears that the use of a task relying on subjective experience can bring out differences in memory and imagery performance.

It could be argued that, in the Imagery condition, participants had no strong motivation to form an accurate mental image, leading to lower contrast accuracy. However, there are number of issues that argue against this possibility. Firstly, participants were clearly instructed that the mental image should correspond to the visual cue, and the accuracy of this would be assessed at the end of the trial. Secondly, orientation discrimination performance was similar in the Imagery and Memory conditions, indicating that participants did attend to the visual cue in the two conditions to a similar extent, and that in both cases, the stimulus was initially encoded equally well. That orientation discrimination performance was similar across Imagery and Memory cases also suggests that the effect of imagery was selective to the feature of the stimulus that participants were encouraged to imagine (i.e. contrast), hence indicating participants engaged in visual imagery or not as they were instructed. Thirdly, no main effect of delay period duration (as well as no interaction between delay duration with the other factors) was observed, indicating that the ability to maintain the mental image did not decay within the time frame (up to 7 seconds) used here.

Moreover, one might argue that participants used verbal codes to hold the contrast and orientation in memory. For example, upon seeing the cue, the participant might label its contrast as a number from 1-6, and remembers that number. We think it is unlikely that such verbal code could be done only in a few trials. There were 6 different contrasts used, and 8 different orientations. Forming verbal labels of these would take considerable number of trials, as realistically, participants would need to see each stimulus type a number of times before realizing how many different stimuli there are, and how they relate to each other. This is not trivial, especially as the visual cues were shown in randomized order. Thirdly, in the debriefing, the use of strategies was discussed with the participants, and all informed having used the given instructions. Thus it is unlikely that verbal recoding can explain this pattern of results.

Interestingly, even when trials requiring the report of “imagery content” or “memory content” were intermixed (in Experiment 2), the difference in accuracy remained.

Note that in Experiment 2 participants did not know at the beginning of each trial whether Imagery or Memory contrast would be assessed, hence they needed to form a representation of the visual cue on each trial to be used in the subsequent matching test. The accuracy difference here indicates that information is accessed differently when the matching judgments is based on memory or imagery, of which the one reflecting memory content leads to improved performance in the matching test. In other words, imagery does not appear to operate on the same representation as the perceptual memory of the visual cue, as the former leads to significantly lower accuracy. This finding suggests that perceptual memory and imagery may not fully share the same neuronal representation, challenging the current view of a close functional and neuroanatomical linkage between memory and imagery (Serences et al., 2009; Harrison & Tong 2009, Albers et al., 2013). Nevertheless, the results of this study add on to existing evidence of that memory and imagery can at least be partly dissociable (e.g; Saad et al., 2015, Saad & Silvanto, 2013; Zeman et al., 2010).

We propose that the finding that overall fidelity of contrast matching is better in the mere memory condition than in the imagery condition may rely on an implicit memory system (see e.g. (Magnussen, 2000; Vogt & Magnussen, 2007) rather than on an explicit system linked to conscious awareness –i.e. during imagery condition. Phenomenal experience of informational content may be a key mental process that distinguishes imagery from other forms of memory (Pearson & Kosslyn, 2015; Pearson et al., 2015). Even some forms of working memory processing (e.g. for maintaining orientation features across delays) may operate on non-conscious input (Soto et al. 2011; Soto & Silvanto, 2014). However we speculate that the act of phenomenally experiencing a cue in the focus of the mind’
ss eye (i.e. imagery) comes at a cost in the fidelity of its representation, which by default, is encoded in a high-resolution primitive perceptual memory system (Magnussen, 2000). This suggestion is in keeping with the observation that high imagers under certain circumstances have lower memory retrieval performance than low imagers (Reisberg et al., 1986; Heuer et al., 1986; Reisberg & Leak, 1987). In conclusion, our results show that using mental imagery, as recall strategy is not helpful to get the most accurate performance. Therefore, these results raise the interesting possibility that visual short-term memory and visual imagery might be based on distinct neural representations (Jacobs & Silvanto, 2015; Saad et al, 2015).

## Notes

### Competing Interest Statement

The authors have declared no competing interest.

